# Brain B Vitamin Status and One-Carbon Metabolism Gene Polymorphisms are Associated with Cognitive Impairment in Alzheimer’s and Parkinson’s Disease

**DOI:** 10.64898/2026.04.22.719490

**Authors:** Karel Kalecký, Jessica Castañeda-Gill, Sharlin Patel, Teodoro Bottiglieri

## Abstract

**Introduction:** Disturbances in one-carbon metabolism and homocysteine (Hcy) regulation have been implicated in Alzheimer’s disease (AD) and Parkinson’s disease (PD), yet direct evidence from human brain tissue and the contribution of genetic variation remain limited. We investigated whether B-vitamin-related metabolic deficits and polymorphisms in one-carbon metabolism pathways contribute to cognitive impairment in AD and PD.

**Methods:** We quantitated metabolites related to B vitamins and one-carbon metabolism in post-mortem frontal cortex (n=136) and putamen (n=68) from clinically and neuropathologically characterized healthy controls, AD dementia (AD-D), PD with dementia (PD-D), PD with mild cognitive impairment (PD-MCI), and cognitively normal PD (PD-CN). Genotyping targeted variants related to one-carbon metabolism and B vitamins. Multivariable regression models controlled for demographic, clinical, metabolic, and tissue-handling covariates.

**Results:** AD-D and PD-D shared a convergent cortical signature of reduced biotin, pyridoxal-5’-phosphate, thiamine monophosphate, pantothenic acid, and betaine. Elevated Hcy and reduced tetrahydrofolate were specific to AD-D and to PD-D during acute levodopa exposure. PD-MCI exhibited deficits in putamen resembling cortical dementia-associated changes. Self-reported B-vitamin supplementation attenuated several abnormalities, including complete normalization of levodopa-associated Hcy and folate disturbances in PD-D. Across subjects, multiple B vitamins showed strong associations with Hcy and betaine, indicating coordinated impairment of all three Hcy-metabolizing pathways. Genotyping analysis identified polymorphisms in MTRR, TYMS, MTR, NFE2L2, BHMT, and MTHFR that were differentially distributed across cognitive subgroups or associated with AD-related pathology scores. Composite genetic and metabolic burden scores were elevated in cognitively impaired groups but not in cognitively normal groups.

**Conclusion:** Dementia in AD and PD is characterized by convergent B-vitamin deficiencies and genetic susceptibilities that disrupt Hcy metabolism. These findings provide a mechanistic explanation for levodopa-related Hcy accumulation in vulnerable PD subjects and identify B-vitamin supplementation as a potentially modifiable factor relevant to cognitive decline.

## Introduction

Alzheimer’s disease (AD) and Parkinson’s disease (PD) are two most prevalent forms of neurodegeneration. While the primary affected brain regions and main symptoms are different, both diseases share several underlying characteristics, including progressive spread of protein aggregates and involvement of oxidative stress, pro-inflammatory environment, and mitochondrial dysfunction^1,2^. Another similarity lies in the risk of developing dementia in PD, which is up to 6 times higher than for others^3^. The histopathology of PD dementia (PD-D), however, differs from AD dementia (AD-D).

Recently, we have reported disturbances in one-carbon metabolism in brain cortex in both AD-D and PD-D^4^. The central point of one-carbon metabolism pathologies is homocysteine (Hcy), a toxic intermediate in the methionine cycle. Elevated plasma total Hcy is a well-established risk factor for development of dementia and AD^5^. Levodopa, a parkinsonian medication with L-3,4-dihydroxyphenylalanine (DOPA), is partially metabolized by catechol-O-methyltransferase (COMT), generating reaction by-products S-adenosylhomocysteine (SAH) and Hcy. Our study showed accumulation of Hcy in brain during acute levodopa presence in PD-D, but not PD without dementia (PD-ND), and elevation of Hcy metabolic products in AD^4^. Moreover, we saw reduced Hcy re-methylation potential in demented subjects, including low betaine (trimethylglycine), which was significantly correlated^4^ with Mini-Mental State Examination (MMSE) cognitive score^6^.

One-carbon metabolism relies on active forms of several B vitamins as cofactors for key enzymes, especially pyridoxal 5’-phosphate (PLP; B6), methyltetrahydrofolate (MTHF; B9), and methylcobalamin (MeCbl; B12). Hcy is a branch-point metabolite that can be converted to cystathionine by PLP-dependent cystathionine β-synthase (CBS), or re-methylated back into methionine by MTHF and MeCbl-dependent methionine synthase (MTR) or betaine-homocysteine methyltransferase (BHMT). There are reports of lower plasma PLP in both AD^7^ and PD^8^. There are also known single nucleotide polymorphisms (SNPs) which affect this pathway, often investigated in the context of neural tube defects^9,10^. Gene variant rs1801133 in methylenetetrahydrofolate reductase (MTHFR) has been associated with faster brain atrophy^11^.

However, there is insufficient research on the levels of B vitamins directly in human brain tissue in relation to neurodegenerative disorders or cognitive impairment. There are only reports of decreased levels of pantothenic acid (B5) in AD^12^ and PD^13^. Our recent metabolomic study in PD brain^14^ found indirect evidence pointing towards lower B5 and PLP.

In the current case-control study, we measured a panel of B vitamin-related compounds in human brain frontal cortex, resp. putamen, and searched for associations with AD and PD in relation to cognitive impairment and dementia. We also genotyped the samples and analyzed the effects of variations in genes involved in one-carbon metabolism and the affected B vitamins. We then integrated the data with our previous measurements of Hcy and betaine and attempted to explain the disturbances observed in one-carbon metabolism in AD and PD^4^.

## Methods

### Study subjects

Post-mortem brain frontal cortex samples (Brodmann area 9, 500 mg) were obtained from the Banner Sun Health Research Institute brain bank (Sun City, AZ, USA)^15^. This included 136 subjects comprised of 36 cognitively normal healthy controls (HC-CN), 35 AD-D subjects, and 65 PD subjects at various stages of cognitive impairment: 32 PD-D and 33 PD-ND cases, subdivided into 19 with mild cognitive impairment (MCI; PD-MCI) and 14 cognitively normal (PD-CN). We also obtained post-mortem putamen samples (20 mg) of the identical 35 HC-CN (1 sample was destroyed during analysis) and 33 PD-ND subjects. The numbers were based on available tissue in the biobank with required characteristics and relatively short post-mortem collection interval range, combined with allocated funding, and exceed the size of most studies with human brain tissue. The samples were collected between 2004 and 2018 from deceased donors and continuously stored at −80 °C.

The diagnosis followed clinical records and histopathological examination: PD subjects had two of the three cardinal clinical signs of resting tremor, muscular rigidity, and bradykinesia, along with pigmented neuron loss and presence of Lewy bodies in substantia nigra, and were treated with levodopa-carbidopa medication. The status of dementia and MCI corresponds to clinical diagnosis. To differentiate from Alzheimer’s type of dementia, the PD-D subjects were allowed to score no more than “low likelihood of AD” in NIA-Reagan classification^16^, whereas AD subjects were classified as “high likelihood of AD”. Controls were without a history of cognitive impairment and parkinsonism. All subjects were White Americans, either non-Hispanic or with ethnicity not provided. No other major pathologies of central nervous system were present.

Clinical profiles with assessment of progression scores were updated during the last ante-mortem visit. Histopathological density scores of neurofibrillary tangles and senile plaque (neuritic, cored, and diffuse) were evaluated using templates of the Consortium to Establish a Registry for Alzheimer’s Disease (CERAD)^17^ and the scores were summed over five brain regions: frontal, temporal, parietal, entorhinal, and hippocampal CA1 region. Other scores of disease progression included MMSE, motor section score of the Unified Parkinson’s Disease Rating Scale (UPDRS-M)^18^, Unified Staging System for Lewy Body Disorders (USSLB)^19^, and disease duration since diagnosis.

### Chromatography and mass spectrometry

We performed targeted triple quadrupole liquid chromatography tandem mass spectrometry analysis integrating 5 separate assays. B vitamins (except for folates) were measured according to transitions from Xu et al.^20^ and Meisser Redeuil et al.^21^, preferring those with stronger signal: the former one for MeCbl, niacin, riboflavin, and thiamine pyrophosphate (TPP), the latter one for biotin, nicotinamide, pantothenic acid, PLP, thiamine, and thiamine monophosphate (TMP). Brain tissue was prepared by extraction using ultrasonic homogenization with ethanol in phosphate buffered saline at the ratio of 85:15 (Et_85_PBS_15_) and tissue concentration 4 µl/mg. Methylmalonic acid (MMA) was quantitated using an approach of Lai et al.^22^ after extraction with 0.1M perchloric acid (3 µl/mg). Measurements and methods for non-protein-bound Hcy and DOPA as well as cortex betaine, MTHF, and tetrahydrofolate (THF) were presented in our previous publications^4,14^ and here we repurposed the data. Due to limited putamen tissue amount, the putamen samples were extracted with Et_85_PBS_15_ (3 µl/mg) and aliquots were dried down, stored in freezer at −80 °C, and later reconstituted to the appropriate volume before usage. They were not subjected to the folate assay, which would require different processing. B-vitamins were separated on a chromatographic column, Phenomenex Luna C18 (Phenomenex, Torrance, California, USA) and all assays were run on Shimadzu Nexera chromatography platform (Shimadzu Corporation, Kyoto, Japan) coupled to Sciex QTrap 5500 mass spectrometer (AB Sciex LLC, Framingham, Massachusetts, USA), except for the folates, which were analyzed using a Waters Xevo TQ MS spectrometer (Waters Corporation, Milford, Massachusetts, USA).

Sample handling was done on dry ice to avoid multiple freeze-thaw cycles. We randomized the samples across plates, with stratification, already prior to their processing to avoid any accidental bias towards one of the subject groups. Plates included blanks to calculate limits of detection, repeats of a quality control sample (QC) to monitor the variation, and at least seven calibrators for all compounds except for several B vitamins in cortex and B vitamins in putamen. Internal standards were not available for the B vitamin assay, the performance of which, however, was very consistent, typically achieving the coefficient of variation (CV) of QC < 5%. Chromatographic peaks were manually reviewed and integrated in the in-house software Integrator developed for high-consistency multi-plate peak integration (patent pending: Application # PCT/US24/51426, filed on October 15, 2024).

### Genotyping

The samples were analyzed on the Illumina iScan platform (Illumina Inc, San Diego, California, USA) with a bead-based microarray Illumina OmniExpress-24 v1.3 and processed in Illumina GenomeStudio Genotyping v2.0.5. The guanine-cytosine content cut-off for successful variant identification was set to 0.15. This assay was selected for its coverage of the main SNPs of interest, combined with its broader scope for the purpose of additional exploration.

### Data analysis

#### Data preprocessing

All data manipulation and analysis was done with R v4.4.2^23^ in RStudio v2024.12^24^. The chromatographic peak areas, in ratio with internal standard peak areas, were calibrated with a fitted quadratic calibration curves where available. Limits of detection (LODs) were calculated as an estimated 99.9th normal distribution percentile (mean + 3.09×standard deviation (SD)) of the signal in blanks in each plate. Only methylcobalamin in cortex was consistently (in majority of samples) below the LOD and was excluded from the analysis. To ameliorate batch effects, multi-plate data were normalized (per metabolite) with a sample distribution-based normalization that respects subject group assignments. To better approximate Gaussian distributions, we applied Box-Cox transformation on the concentration values with R package *car*^25^. Remote outliers were detected and adjusted with conventional Tukey’s fencing^26^ (using k = 3) to protect against skewing the means by extreme values while not reducing the variance greatly. Finally, the values were standardized with respect to control samples to facilitate comparison of regression coefficients in the statistical analysis. In the genomic data, rare SNPs with the alternate allele present in less than 5 samples were filtered out. Since regression requires all covariates to be non-missing, 4 missing body mass index (BMI) values and 6 missing education values were imputed as a mean value conditional on the subject group and sex. Among measured values, we found no THF peak in 4 samples and interpolated these values as half of the least detected THF concentration.

#### Sociodemographic and clinical characteristics

Key characteristics of subjects and samples in each group were compared with Fisher’s exact test (binomial variables) and analysis of variance (ANOVA) on a linear model constructed with the R package *nlme*^27^ (continuous variables), allowing for group-specific variance.

#### Metabolomic data

The differential analysis between controls and other subject groups was based on a series of multivariable linear regression models with R package *nlme*^27^, one for each metabolite, with its values as a dependent variable and subject groups – either separately, or with PD subgroups or demented subgroups together – as independent variables, allowing for group-specific variance. The models included covariates for age, sex, education, BMI, acute levodopa medication presence at the time of death (estimated from log-transformed DOPA concentrations as values in PD subjects above the estimated normal distribution 95th percentile of the values of controls in the respective brain region; separate covariates for PD-D and PD-ND following the DOPA interaction^14^), diagnosis of several frequent disorders affecting metabolism: hyperlipidemia, diabetes mellitus, renal insufficiency, and hypothyroidism, as well as post-mortem collection interval and the total length of freezer storage. The last two covariates were log-transformed to be able to capture any time-related exponential decay. Due to standardization, the regression coefficients have a unit of 1 standard deviation of the distribution of controls. Assessment of collinearity among all regressors was based on the magnitude of Pearson’s correlation coefficients and adjusted generalized variable inflation factor (GVIF) calculated with the R package *car*^25^. We found no evidence of significant collinearity (all Pearson’s |r| < 0.6 and adjusted GVIFs < 1.5). The analysis of associations with disease progression scores followed the same regression model, with the vitamins B-related compounds (one at a time) included as an additional independent variable and the progression score being the dependent variable, and was also done separately in cognitive and diagnostic subgroups. The analysis of associations with Hcy and betaine followed the same regression model, with these compounds being the dependent variables, and was done in the whole cohort to search for general associations. We kept all covariates, including group assignments, to account for their impact on B vitamins levels.

#### Genomic data

We first focused on 9 pre-selected SNPs. In the second round, we included all available non-synonymous SNPs related to metabolism of homocysteine, betaine, and B vitamin-related compounds that we found altered in the subject groups. The differential presence of SNPs among the subject groups was compared with a series of Wilcoxon-Mann-Whitney tests with R package *coin*^28^, also done separately in cognitive and diagnostic subgroups. The association analysis with disease progression scores was performed with a series of linear regression models analogically to metabolomic data, except for exclusion of covariates relevant only to physical samples (post-mortem collection interval, length of freezer storage, and levodopa presence in tissue). For the association analysis with Hcy and betaine and the later calculated composite score, we first decided for each SNP whether it should be treated as dominant, recessive, or additive by constructing different linear models with one of the three effect types as the independent variable and values of the respective metabolite as the dependent variable, adjusted with other covariates, and compared the models with the Bayesian information criterion. Unlike the analysis with B vitamins, it was not necessary to include all covariates, and we considered only objective and tissue-related covariates (sex, age, post-mortem collection interval, and the total length of freezer storage). However, samples with acute levodopa presence were not considered in this part due to its specific effects on Hcy and betaine. Linkage disequilibrium statistics were calculated with R package *genetics*^29^.

#### False discovery rate (FDR) control

For each regression coefficient of interest, two-tailed p-values across the series of models were controlled for FDR with the Benjamini-Hochberg procedure^30^ or q-value technique with R package qvalue^31^ in case of larger distributions (the extended SNP analysis). Effects with FDR ≤ 0.05 and ≤ 0.10 were considered statistically significant for the metabolomic and genomic analysis, respectively.

#### Composite scores

The general associations with Hcy and betaine were aggregated to calculate 3 composite scores: A) SNP risk score – number of polymorphisms associated with elevated Hcy or decreased betaine (respecting the estimated SNP model – recessive, dominant or additive), B) SNP protection score – number of polymorphisms associated with decreased Hcy or increased betaine, and C) B vitamins risk score – number of B vitamins negatively associated with Hcy or positively associated with betaine with the level below the estimated 5th percentile (z-score −1.64) of the normalized distribution of controls. The statistical evaluation was done as pair-wise group comparisons with HC-CN using Wilcoxon-Mann-Whitney tests with R package *coin*^28^.

#### Combinatorial analysis

The interactions between groups, metabolites, and SNPs were further evaluated with a genetic programming approach. We implemented the algorithm in C# in Microsoft Visual Studio 2022 (Microsoft Corporation, Redmond, Washington, USA) with the typical set of features including combination, mutation, and cross-over in the population of evolving expression trees. Operands consisted of Boolean expressions from values of metabolites with a threshold in terms of standard deviations from the mean of controls (−2, 1, 0, 1, 2) and SNPs combined with a threshold on the number of affected alleles (0, 1, 2), or respectively, as numeric expressions of the number of affected alleles. Operators were both logical (negation, logical OR, logical AND) and arithmetic (add, subtract, multiply, negative). The fitness score was calculated as an average of classification accuracies for both subgroups of interest, penalized 5 percent points (p.p.) for every operand in the expression tree. We also incorporated a noise term which gradually decreased from up to 10 p.p. (following a uniform random distribution) to 0 during the first half of the run to mimic the simulated annealing technique. The population size was set to 100,000 with 100 evolution steps. The best result of 10 repeated runs was selected. Primarily, we performed this analysis as explorative although we also provide 10-fold cross-validation evaluation.

## Results

### Cohort overview

Basic sociodemographic and clinical characteristics of the subjects are summarized in **Table 1**. Sociodemographic variables show no significant differences between the groups. Levodopa medication dosage was similar across the PD groups although this information is available only for a third of the subjects. Half of the AD-D subjects had apolipoprotein E (APOE) ε4 allele (ε4+) in concordance with its relevancy for the disease. Scores of disease progression and histopathological examination showed major differences as expected. There were no differences in several disorders that impact metabolism, specifically hyperlipidemia, diabetes mellitus, renal insufficiency, and hypothyroidism.

**Table 1.** Sociodemographic and clinical characteristics of the subjects.

|  | Groups |  |  |  |  | P-value |
| --- | --- | --- | --- | --- | --- | --- |
|  | Controls<br>(N = 36) | PD-CN<br>(N = 14) | PD-MCI<br>(N = 19) | PD-D<br>(N = 32) | AD<br>(N = 35) |  |
| Sex |  |  |  |  |  |  |
| Male, No. (%) | 22 (61) | 8 (57) | 14 (74) | 24 (75) | 20 (57) | 0.47 |
| Female, No. (%) | 14 (39) | 6 (43) | 5 (26) | 8 (25) | 15 (43) |  |
| Ethnicity |  |  |  |  |  |  |
| Non-Hispanic, No. (%) | 32 (89) | 14 (100) | 16 (84) | 28 (88) | 34 (97) | 0.29 |
| Not provided, No. (%) | 4 (11) | 0 (0) | 3 (16) | 4 (13) | 1 (3) |  |
| Race |  |  | All White |  |  | 1 |
| Age, years, mean (SD) | 82 (10) | 85 (6) | 82 (6) | 80 (5) | 81 (9) | 0.10 |
| Education, years, mean (SD) | 14 (3) | 16 (3) | 14 (3) | 16 (3) | 15 (2) | 0.05 |
| BMI, kg/m2, mean (SD) | 25 (5) | 22 (4) | 25 (8) | 27 (9) | 24 (4) | 0.08 |
| Levodopa dosage, mg/day, mean (SD) | - | 450 (217) | 500 (173) | 509 (226) | - | 0.86 |
| APOE ε4, allele presence, No. (%) | 5 (14) | 2 (14) | 3 (16) | 3 (9) | 18 (51) | <0.001 |
| Disease duration, years, mean (SD) | - | 11 (7) | 12 (6) | 18 (9) | 9 (4) | <0.001 |
| UPDRS-M score, mean (SD) | 7 (5) | 32 (15) | 31 (15) | 50 (16) | 17 (18) | <0.001 |
| USSLB score, mean (SD) | 0.0 (0.0) | 3.0 (0.5) | 3.7 (0.5) | 3.4 (0.6) | 0.0 (0.0) | <0.001 |
| MMSE score, mean (SD) | 28 (1) | 28 (1) | 25 (3) | 20 (6) | 15 (8) | <0.001 |
| Neurofibrillary tangle density, mean (SD) | 2.9 (1.6) | 5.3 (2.6) | 5.2 (1.6) | 4.8 (2.1) | 14 (1.7) | <0.001 |
| Senile plaque density, mean (SD) | 2.7 (3.9) | 4.6 (4.7) | 5.1 (6.0) | 1.8 (3.3) | 14 (1.0) | <0.001 |
| Freezer storage, years, mean (SD) | 12 (4) | 10 (4) | 8 (4) | 10 (4) | 9 (2) | <0.001 |
| PMCI, hours, mean (SD) | 3.2 (1.0) | 3.3 (0.9) | 3.4 (1.2) | 3.4 (0.9) | 3.4 (0.8) | 0.87 |
| Hyperlipidemia, No. (%) | 13 (36) | 7 (50) | 9 (47) | 11 (34) | 20 (57) | 0.30 |
| Diabetes mellitus, No. (%) | 12 (33) | 1 (7) | 3 (16) | 8 (25) | 6 (17) | 0.27 |
| Renal insufficiency, No. (%) | 5 (14) | 3 (21) | 2 (11) | 5 (16) | 2 (6) | 0.53 |
| Hypothyroidism, No. (%) | 9 (25) | 3 (21) | 5 (26) | 5 (16) | 3 (9) | 0.32 |

Levodopa dosage is unambiguously known only for 31% of PD subjects. MMSE score is known for 87% subjects. UPDRS-M score, measured in off-levodopa state, is known for 57% of PD subjects and 59% others.
Abbreviations: AD, Alzheimer's disease; APOE, apolipoprotein E; BMI, body-mass index; CN, cognitively normal; D, dementia; MCI, mild cognitive impairment; MMSE, Mini-Mental State Examination; PD, Parkinson's disease; PMCI, post-mortem collection interval; UPDRS-M, motor section of the Unified Parkinson's Disease Rating Scale; USSLB, Unified Staging System for Lewy Body Disorders.

Importantly, post-mortem collection intervals were very short and homogeneous between the groups. Duration of freezer storage was somewhat higher for controls, but we did not observe any signs of related tissue degradation (e.g. choline levels of the longer stored control samples were indistinguishable from those with shorter storage; Welch’s t-test p-value = 0.72). Any potential freezer storage effect was also controlled for in the regression analysis alongside other covariates.

### AD-D and PD-D show similar B vitamins alterations in brain cortex

First, we focused on differential analysis of B vitamins and related compounds among subject groups (**Figure 1**). Besides increased Hcy in AD-D and decreased betaine in both AD-D and PD-D from previously reported data^4^, we found lower cortical levels of biotin (normalized regression coefficient (β) = −0.73, p = 0.003, FDR = 0.023), THF (β = −0.53, p = 0.012, FDR = 0.043), and PLP (β = −0.51, p = 0.017, FDR = 0.049) in AD-D. Biotin and PLP showed a similarly strong trend in PD-D and both compounds increased their significance when considering all demented subjects together, additionally revealing decreased TMP (β = −0.51, p = 0.009, FDR = 0.032) and pantothenic acid (β = −0.64, p = 0.010, FDR = 0.032). Thus, 5 changes constitute the dementia-related association: decreased betaine, biotin, PLP, TMP, and pantothenic acid. On the contrary, increased Hcy and decreased THF were specific to AD-D with no similar trends in PD-D. However, when analyzing the effect of acute levodopa presence, we saw the same change in Hcy (as previously reported^4^) as well as THF, both occurring only in PD-D with levodopa presence (L+) while PD-ND samples remained unchanged, and both changes reached roughly double the magnitude compared to AD-D.

**Fig. 1:**
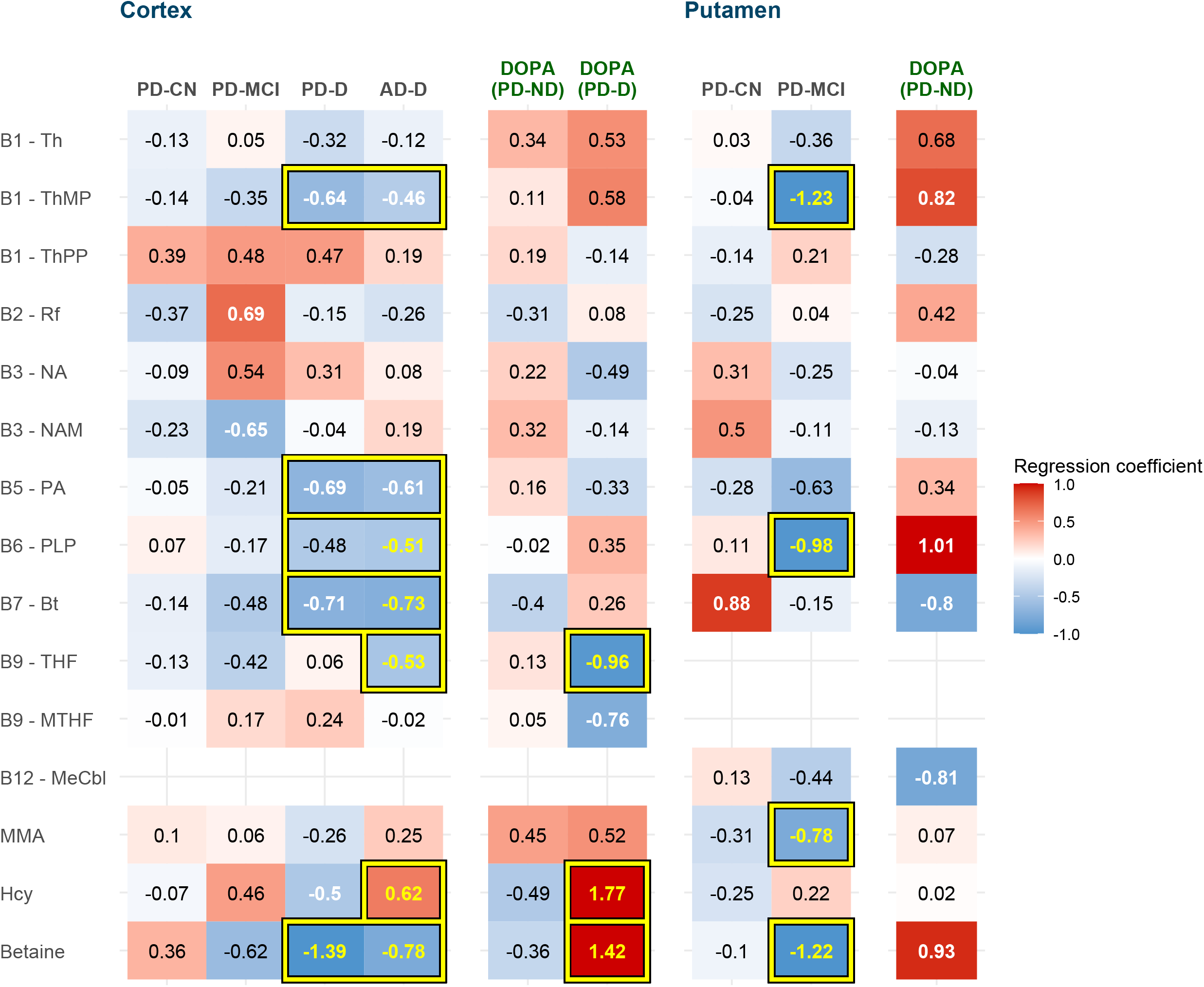
Metabolite associations with Groups (with DOPA effect) Group heatmap with disturbances in B vitamin-related compounds: Heatmap of normalized regression coefficients from regression analysis with adjustment. Color: blue = lower, red = higher (as compared to controls). Statistically significant differences (FDR ≤ 0.05) are highlighted with yellow font color and framed. Cells framed together are significant when considered both groups together. Nominal significance (p ≤ 0.05) is denoted with white font color. Abbreviations: AD, Alzheimer’s disease; Bt, biotin; CN, cognitively normal; D, dementia; DOPA, indicator of acute levodopa presence; Hcy, homocysteine; MCI, mild cognitive impairment; MMA, methylmalonic acid; MTHF, methyltetrahydrofolate; MeCbl, methylcobalamin; NA, nicotinic acid; NAM, nicotinamide; PA, pantothenic acid; PD, Parkinson’s disease; PLP, pyridoxal 5’-phosphate; Rf, riboflavin; Th, thiamine; ThMP, thiamine monophosphate; ThPP, thiamine pyrophosphate; THF, tetrahydrofolate.

In putamen, where we had no demented subjects, we observed multiple changes in PD-MCI resembling those of dementia in cortex (**Figure 1**) – decreased TMP (β = −1.23, p = 0.0002, FDR = 0.001), betaine (β = −1.22, p = 0.0002, FDR = 0.001), and PLP (β = −0.98, p = 0.006, FDR = 0.026). Additionally, MMA was decreased (β = −0.78, p = 0.012, FDR = 0.039).

### B vitamin supplementation prevents several alterations in brain

We then searched for evidence whether supplementing B vitamins could ameliorate the observed changes. Based on self-reported lists of medications and supplements, which may be incomplete or not up to date at the time of death, we directly compared subjects who reported intake of relevant B vitamins or multivitamins with those who did not (**Figure 2**) among individual subject groups. Most importantly, the levodopa-related increase in Hcy in PD-D disappeared with reported supplements (normalized mean (µ) 1.62 → 0.24, p = 0.025), and parallelly, the decreased THF also normalized (µ −1.23 → −0.33, p = 0.030). Among other dementia-related alterations, pantothenic acid and PLP showed consistent correction in individual subgroups with a varied degree of significance, whereas TMP, biotin, and betaine showed little to no improvement. Furthermore, Hcy was the only compound that manifested the change already in controls (cortex: µ 0.40 → −0.45, p = 0.002; putamen: µ 0.13 → −0.21, p = 0.005).

**Fig. 2:**
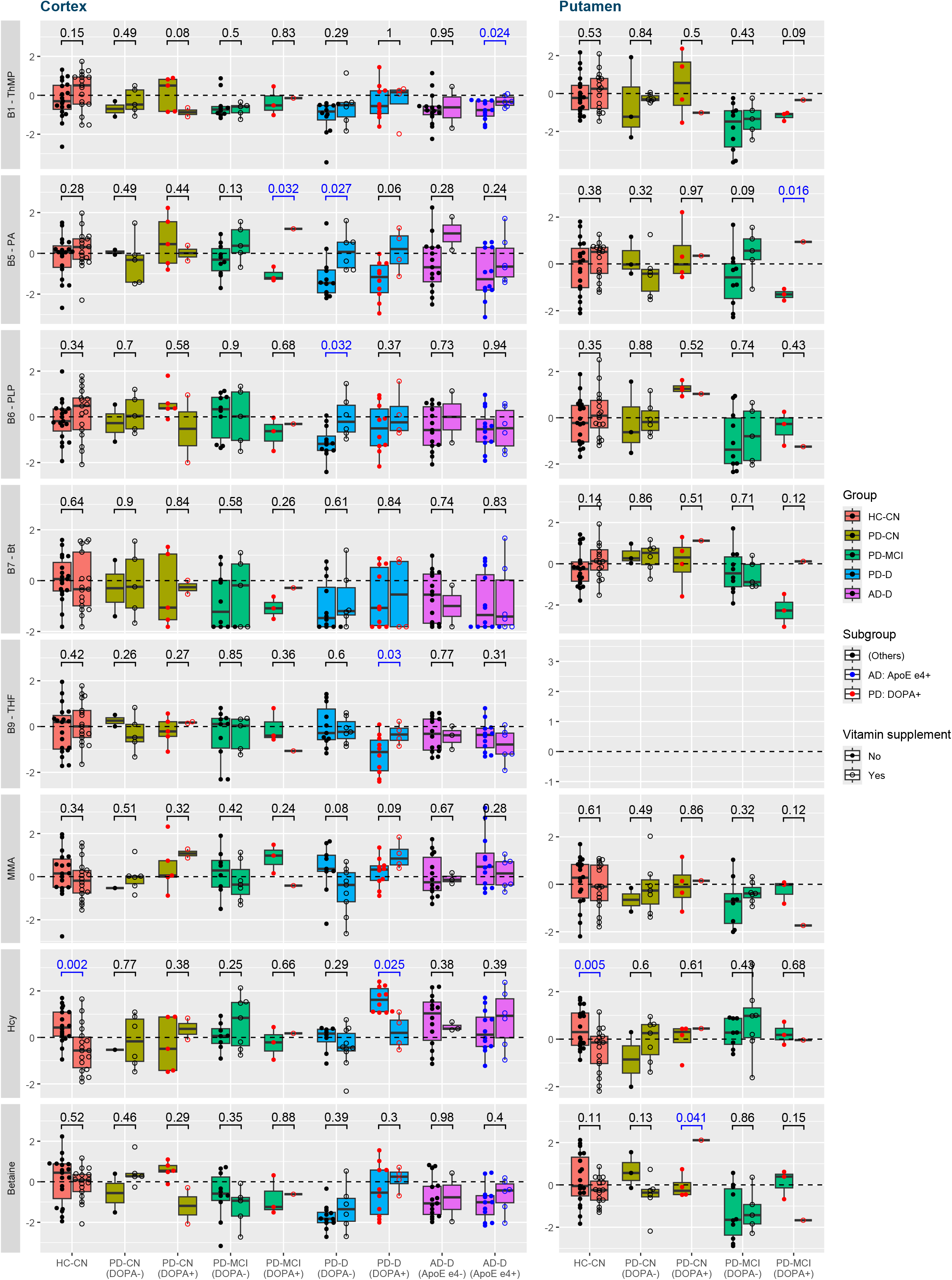
Vitamin supplementation effect in Subgroups. Box plots comparing levels of the altered compounds between subjects who reported the relevant B vitamin supplementation intake and who did not, by subgroup. Differences with nominal significance (p ≤ 0.05) are highlighted in blue font color. Abbreviations: AD, Alzheimer’s disease; Bt, biotin; CN, cognitively normal; D, dementia; DOPA, indicator of acute levodopa presence; HC, healthy control; Hcy, homocysteine; MCI, mild cognitive impairment; MMA, methylmalonic acid; PA, pantothenic acid; PD, Parkinson’s disease; PLP, pyridoxal 5’-phosphate; ThMP, thiamine monophosphate; THF, tetrahydrofolate.

### Levels of multiple B vitamins are associated with changes in Hcy and betaine

Next, we searched for general associations between B vitamins and Hcy and betaine among all subjects together but controlling for the a priori group differences and other covariates. As expected, Hcy was strongly negatively associated with folates (THF: β = −0.42, p = 8e-7, FDR = 1e-5; MTHF: β = −0.30, p = 8e-5, FDR = 0.0005) and PLP (β = −0.29, p = 0.0002, FDR = 0.0007) in cortex (**Figure 3**). Additional relationships include thiamine phosphates (TPP: β = −0.19, p = 0.017, FDR = 0.041; similarly TMP: β = −0.19, p = 0.038, FDR = 0.08) and pantothenic acid (β = −0.16, p = 0.017, FDR = 0.041).

**Fig. 3:**
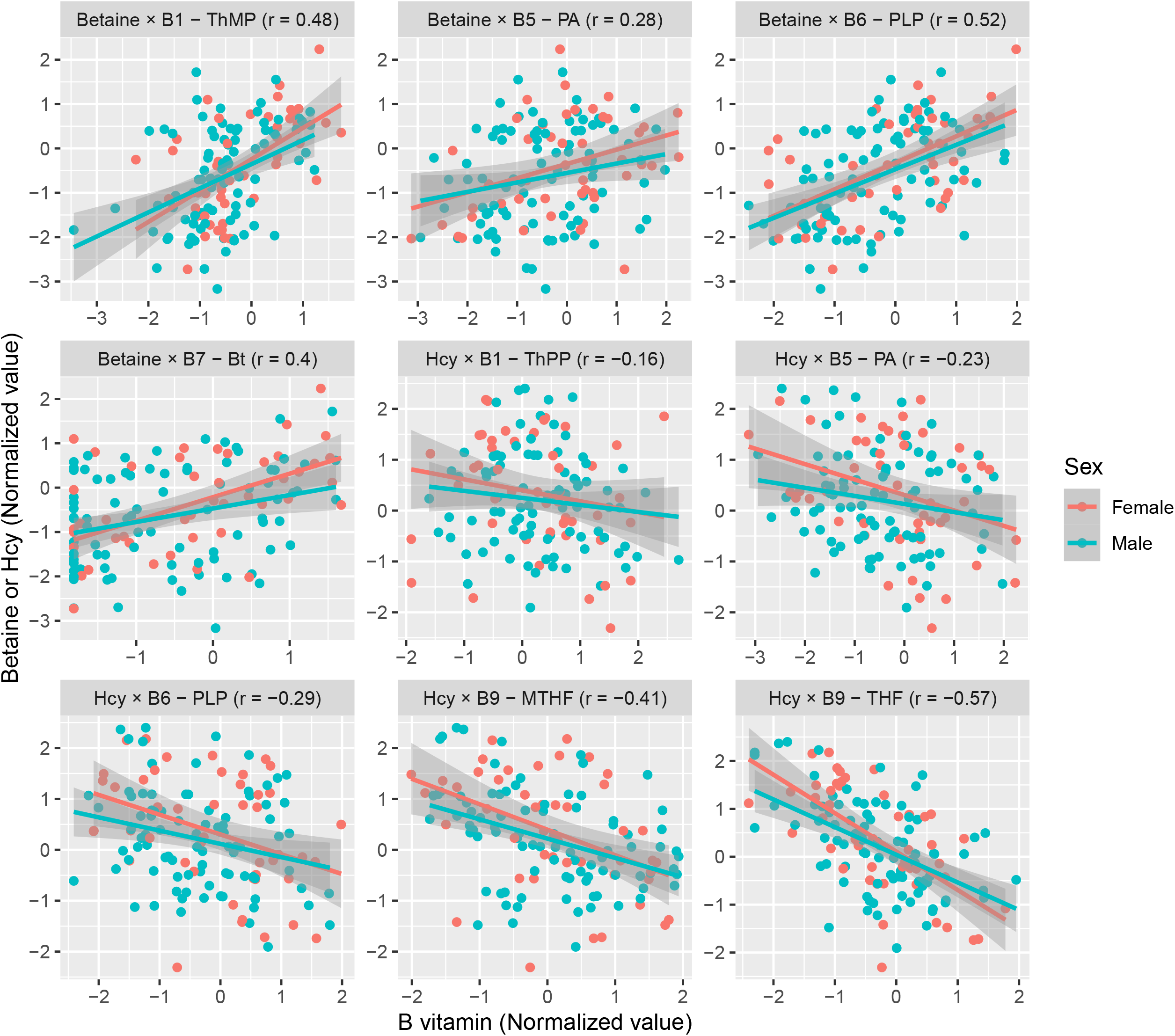
B vitamins associations with Betaine and Hcy. Scatter plots for B vitamins with a significant association (FDR ≤ 0.05) with Hcy or betaine. Linear trend lines are shown by sex to show their consistency. The compounds along with Pearson’s correlation coefficient (r) are specified in subplot headers. Abbreviations: Bt, biotin; Hcy, homocysteine; MTHF, methyltetrahydrofolate; PA, pantothenic acid; PLP, pyridoxal 5’-phosphate; ThMP, thiamine monophosphate; ThPP, thiamine pyrophosphate; THF, tetrahydrofolate.

We saw similar results for betaine in cortex (**Figure 3**), which was positively associated with PLP (β = 0.55, p = 5e-9, FDR = 6e-8), TMP (β = 0.49, p = 1e-5, FDR = 7e-5), pantothenic acid (β = 0.22, p = 0.007, FDR = 0.020), and additionally with biotin (β = 0.31, p = 0.0003, FDR = 0.001). The similarities are despite only a weak association between Hcy and betaine (β = −0.19, p = 0.06).

There were no sex-specific interactions with the associations, with the male and female trend lines overlapping (**Figure 3**). No association was statistically significant in the smaller putamen cohort for both Hcy and betaine although there were identical trends for PLP and pantothenic acid (both Hcy and betaine), and TMP (Hcy only). All results are provided in **Supplementary Table 1**.

### Genetic polymorphisms are associated with cognitive subgroups in PD

In the next step, we performed analysis of SNPs. In each part, we always first focused on 9 pre-selected SNPs, which are directly related to one-carbon metabolism and were previously explored by Wang et al.^9^ and included in the SNP assay: rs234706 (CBS C699T), rs1801133 (MTHFR C677T), rs1801131 (MTHFR A1298C), rs1805087 (MTR A2756G), rs1801394 (methionine synthase reductase (MTRR) A66G), rs1806649 (nuclear factor erythroid-derived 2-like 2 (NFE2L2) C11108T), rs1051266 (reduced folate carrier 1 (SLC19A1) G80A), rs1801198 (transcobalamin 2 (TCN2) G776C), and additionally rs3733890 (BHMT G716A) for its involvement in betaine processing. This was followed by an extended search through all non-synonymous SNPs included in the assay that are related to one-carbon metabolism and the altered B vitamins. For SNPs, we also considered a less stringent FDR threshold (0.1).

We found several associations with cognitive subgroups (**Figure 4A**), notably in PD where we observed significantly lower rate of mutated alleles in PD-D compared to PD-CN for rs2853533 in thymidylate synthase (TYMS; PD-D 6% heterozygous (Het) and 0% homozygous mutant (Hm), PD-CN 50%/7% Het/Hm, p = 0.0001, FDR = 0.006) and rs10380 in MTRR (PD-D 3%/0% Het/Hm, PD-CN 29%/7% Het/Hm, p = 0.003, FDR = 0.058). In both cases, the rate of mutated alleles of PD-MCI subjects were between those of PD-D and PD-CN.

**Fig. 4:**
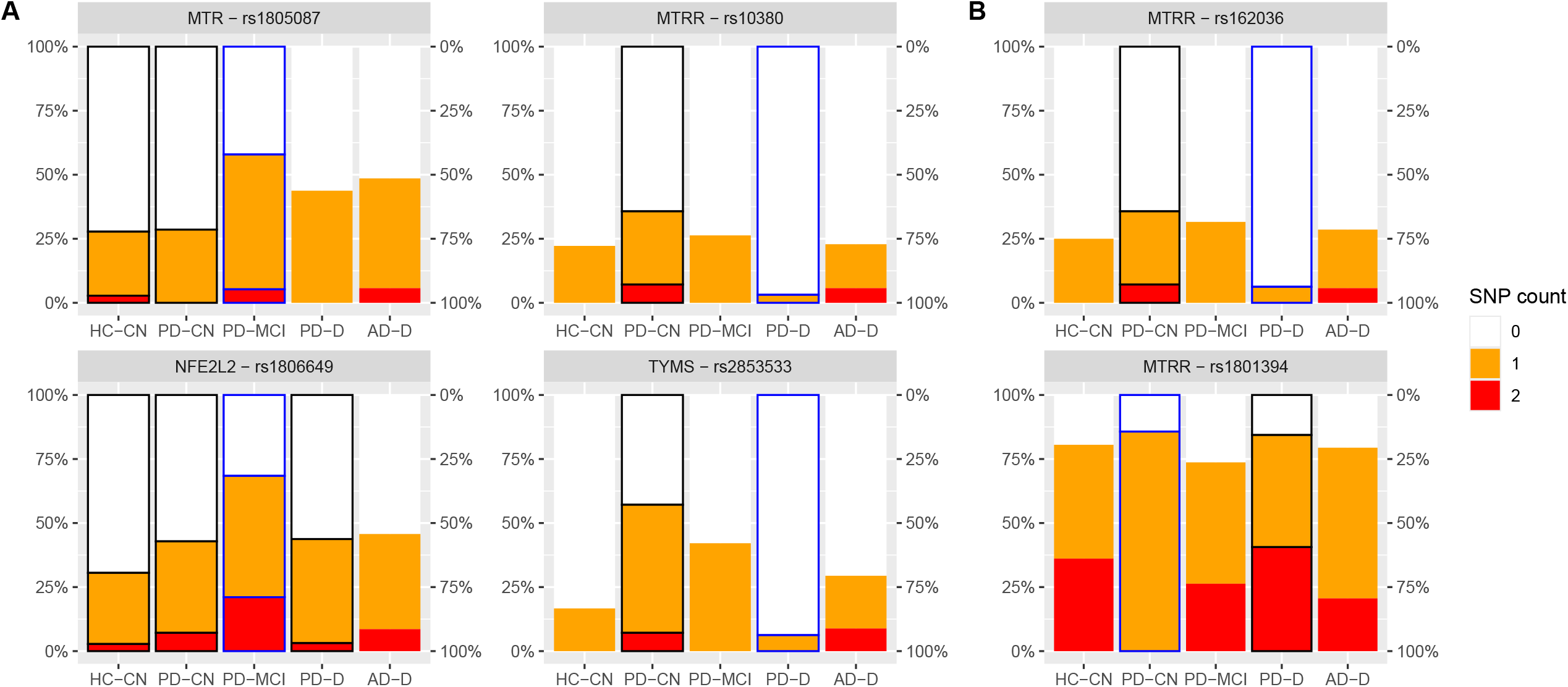
SNP associations with Subgroups. Frequencies of polymorphism among groups: Heatmap representation of frequencies of polymorphisms with (A) significant difference (FDR ≤ 0.1) between groups and (B) additional related polymorphisms with nominal significance (p ≤ 0.05). The differences were significant only when comparing the highlighted group (blue border) with other selected groups (black border). Abbreviations: AD, Alzheimer’s disease; CN, cognitively normal; D, dementia; HC, healthy controls; MCI, mild cognitive impairment; PD, Parkinson’s disease; SNP, single nucleotide polymorphism.

Two other SNPs were significantly associated with PD-MCI subjects (**Figure 4A**): Among non-demented subjects, PD-MCI had a higher rate of mutations compared to PD-CN and HC-CN for rs1805087 in MTR (PD-MCI 53%/5% Het/Hm, PD-CN 29%/0% Het/Hm, HC-CN 25%/3% Het/Hm, p = 0.022, FDR = 0.097). Among all PD subjects and controls, PD-MCI had an increased rate of rs1806649 in NFE2L2 (PD-D 41%/3% Het/Hm, PD-MCI 47%/21% Het/Hm, PD-CN 36%/7% Het/Hm, HC-CN 28%/3% Het/Hm, p = 0.005, FDR = 0.049).

Furthermore, we inspected these 4 genes with associations to see if there were any other related SNPs with similar trends, which could bring more evidence of their functional involvement. The dataset contained only 3 additional SNPs for MTRR but not for other genes. Among these 3 SNPs, 2 reached a nominal significance – rs162036 (PD-D 6%/0% Het/Hm, PD-CN 29%/7% Het/Hm, p = 0.010) and rs1801394 (PD-D 44%/41% Het/Hm, PD-CN 86%/0% Het/Hm, p = 0.044), for which PD-MCI subjects had allele rates also between those of PD-D and PD-CN (**Figure 4B**). While rs162036 and rs10380 were highly correlated (agreement in 96% cases), constituting a single haplotype, rs1801394 was rather anti-correlated, showing mutual exclusivity (with only 13% of Het-Het cases but no Het-Hm or Hm-Hm case; normalized linkage disequilibrium (D’) > 0.99, p = 3e-10).

### Genetic polymorphisms are associated with scores of disease progression

Analysis of SNPs and scores of disease progression found no associations with PD in terms of UPDRS-M or USSLB. However, we discovered several significant associations with markers of AD (**Figure 5**): neurofibrillary tangles increased with the count of rs3733890 (BHMT) mutated alleles (p = 0.002, FDR = 0.020) in AD and PD but not in HC-CN. Senile plaque increased with the count of rs1801131 (MTHFR) mutated alleles and especially in Hm cases (p = 0.002, FDR = 0.015) across the groups except for AD where the plaque score was saturated and uniformly nearing the maximum. On the other hand, rs1801133 (MTHFR) followed the opposite association compared to rs1801131 (p = 0.015, FDR = 0.066). This is likely due to their mutual exclusivity (no Het-Hm and Hm-Hm cases; D’ > 0.99, p = 8e-14), as their combination is frequently non-viable^32^. Additionally, the length of life after diagnosis in subjects with dementia decreased with the presence of rs2294757 in vannin 1 (VNN1), which acts as a pantetheinase and contributes to recycling of pantothenic acid, by 5.1 years in PD-D and 4.6 years in AD-D (p = 0.001, FDR = 0.062). This was regardless of the actual age of death and age of disease diagnosis.

**Fig. 5:**
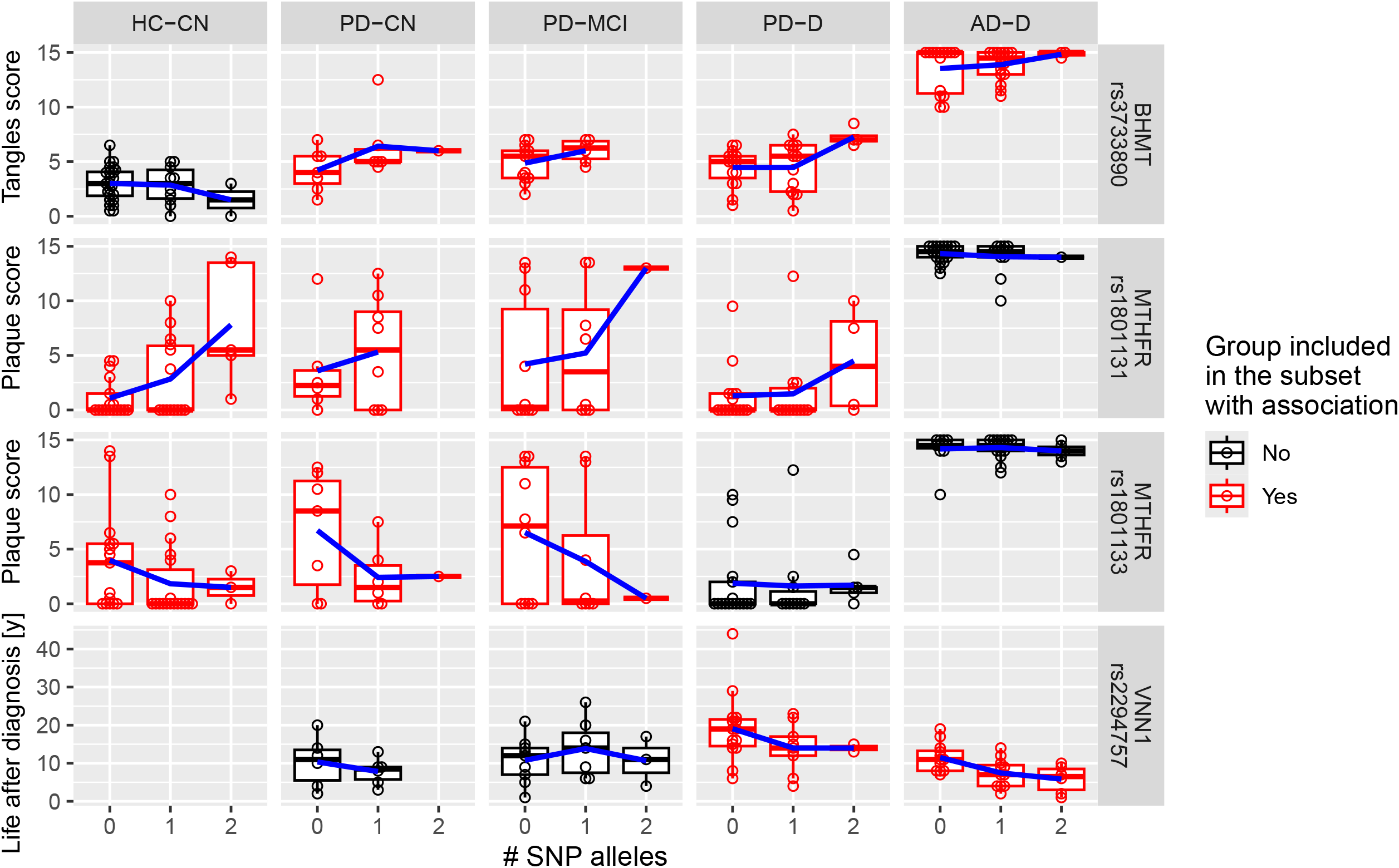
SNP associations with Disease scores. Distribution of disease scores by SNP allele count for associations with significant differences (FDR ≤ 0.1). The associations are valid within the subject groups highlighted in red. The trend lines connect the group means. Abbreviations: AD, Alzheimer’s disease; CN, cognitively normal; D, dementia; HC, healthy controls; Hcy, homocysteine; MCI, mild cognitive impairment; PD, Parkinson’s disease; SNP, single nucleotide polymorphism.

### Genetic polymorphisms are associated with changes in Hcy and betaine

Next, we searched for associations of SNPs with Hcy and betaine and we found two clusters of results (**Figure 6**). The first includes 3 SNPs related to the folate-dependent re-methylation pathway and the mutations had a negative effect on Hcy or betaine: rs1801394 (MTRR) Hm had increased Hcy in putamen (p = 0.0008, FDR = 0.007), rs1805087 (MTR) Het and Hm had decreased betaine in cortex (p = 0.009, FDR = 0.084), and rs2273697 in adenosine triphosphate binding cassette subfamily C member 2 (ABCC2, which facilitates folate transport) Het had decreased betaine in putamen (p = 0.007, FDR = 0.080).

**Fig. 6:**
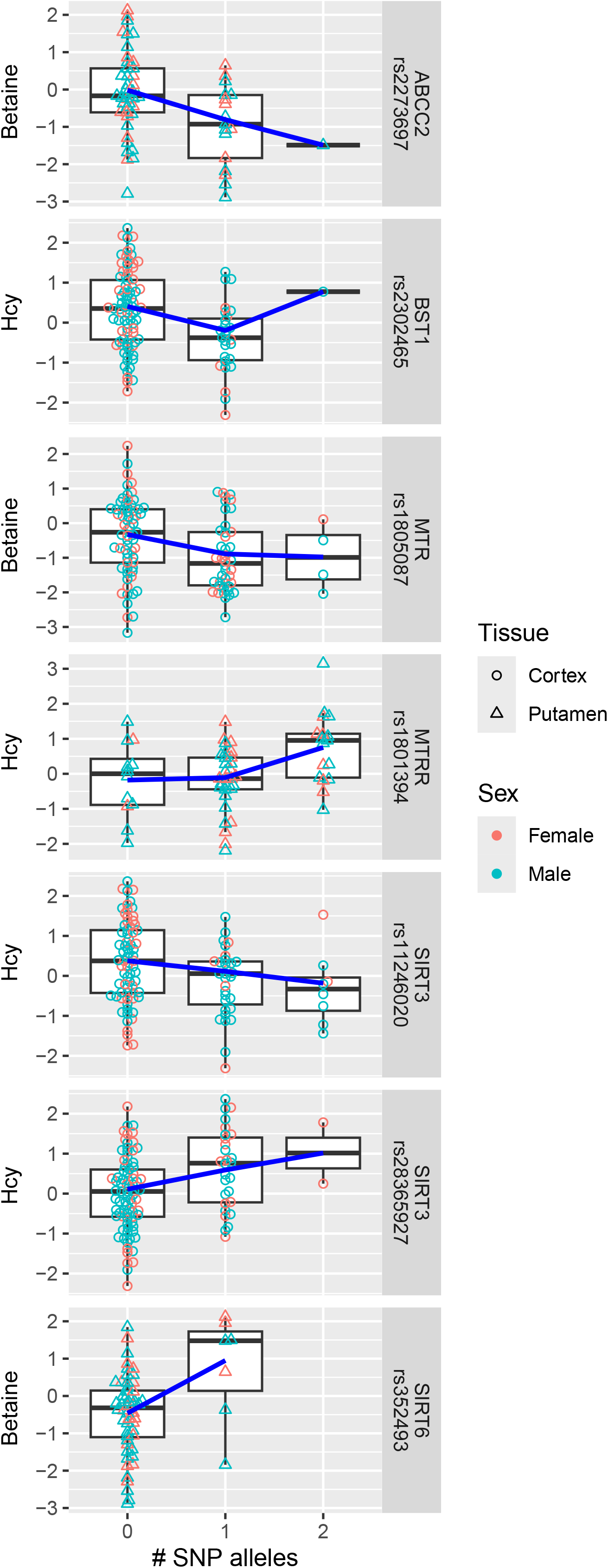
SNP associations with Betaine and Hcy. Polymorphisms associated with Hcy or betaine: Overview of polymorphisms with significant association (FDR ≤ 0.1) in cortex and putamen plotted against transformed Hcy or betaine values across samples. Abbreviations: Hcy, homocysteine; SNP, single nucleotide polymorphism.

The other set of significant associations contains 4 SNPs in genes that convert niacin to nicotinamide and their mutations have mostly a positive effect on Hcy or betaine with the number of mutated alleles (**Figure 6**): rs2302465 in bone marrow stromal cell antigen 1 (BST1) is associated with lower Hcy in cortex (p = 0.003, FDR = 0.040), rs352493 in sirtuin 6 (SIRT6) is associated with increased betaine in putamen (p = 0.002, FDR = 0.046), rs11246020 in sirtuin 3 (SIRT3) is associated with decreased Hcy in cortex (p = 0.006, FDR = 0.055), whereas rs28365927 (SIRT3) was associated in the opposite direction (p = 0.001, FDR = 0.035). These two SIRT3 polymorphisms also show linkage disequilibrium (no Het-Hm and Hm-Hm cases; D’ > 0.99, p = 0.0002).

### Composite scores highlight differences between cognitive subgroups

The detected general associations with Hcy and betaine were further aggregated into several composite scores to see if any subject groups showed a differential burden or protection for increased Hcy or decreased betaine regardless of their actual levels. Interestingly, all cognitively impaired groups exhibited a higher genetic burden (**Figure 7A**): PD-MCI (mean score difference (Δ) = +70%, p = 0.026), PD-D (Δ = +47%, p = 0.040), and AD-D (Δ = +56%, p = 0.037), while PD-CN showed no difference (Δ = +15%, p = 0.58). Within AD-D, the score was more increased in subjects without the APOE ε4 allele (ε4-; Δ = +75%, p = 0.024) than ε4+ subjects (Δ = +38%, p = 0.23). On the other hand, the SNP protection score (**Figure 7B**) was higher in PD-CN (Δ = +40%, p = 0.025), with a trend of decrease in AD-D ε4-(Δ = −31%, p = 0.083) but not AD-D ε4+ (Δ = −13%, p = 0.57).

**Fig. 7:**
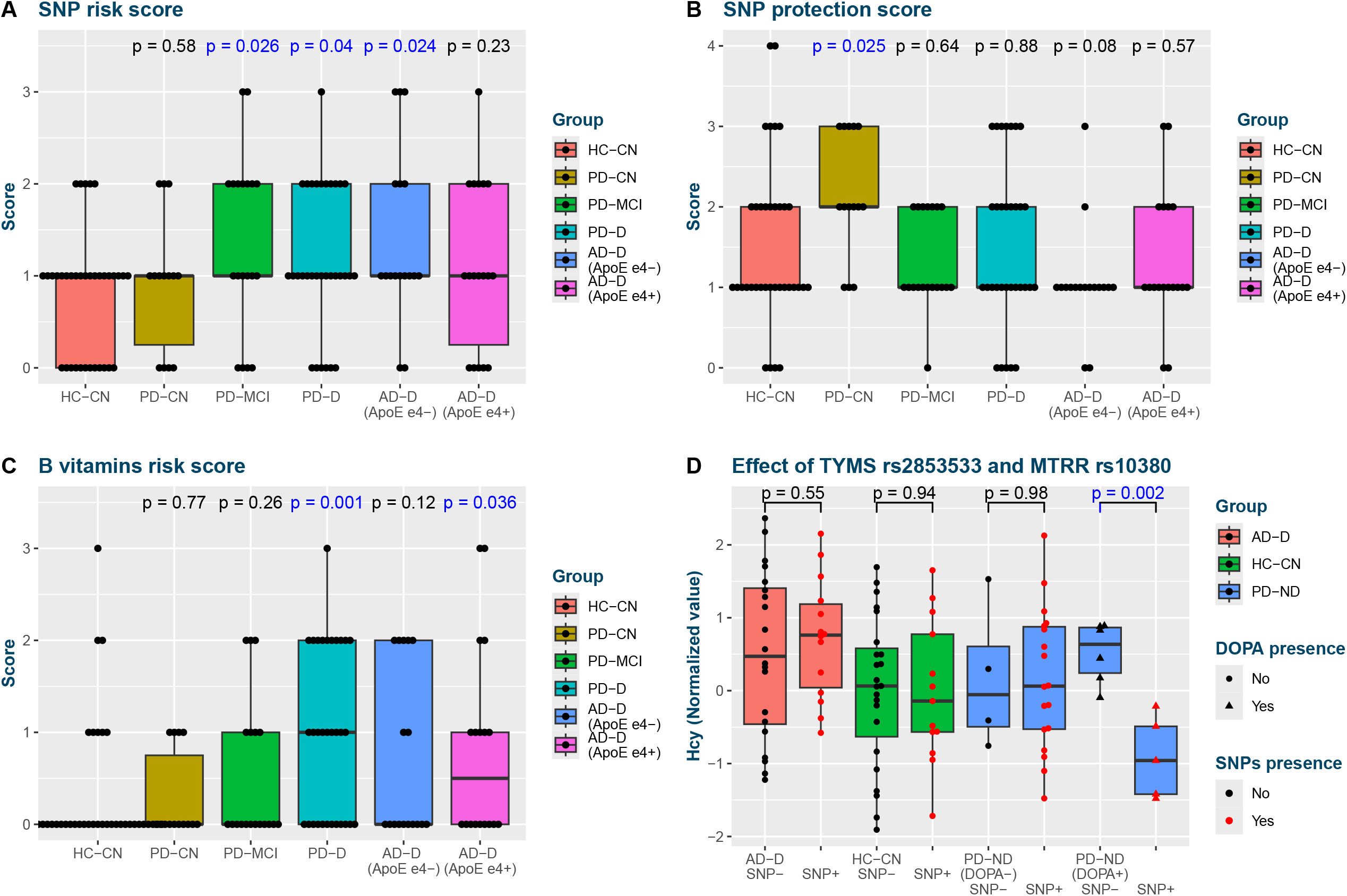
Composite scores. Composite analysis: Group distributions of one-carbon metabolism composite scores based on Hcy and betaine associations as compared with controls: SNP risk score (A), SNP protection score (B), and B vitamins risk score (C). (D) Effect of presence of one of TYMS G5461C and MTRR C1783T polymorphisms on Hcy levels, which is visible only in subjects with acute levodopa presence (L+). Note that this SNP combination was selected to best differentiate PD-D from PD-ND, i.e. independently from the phenomenon observed here. Comparisons with nominal significance (p ≤ 0.05) are highlighted with blue font color. Abbreviations: AD, Alzheimer’s disease; CN, cognitively normal; D, dementia; DOPA, indicator of acute levodopa presence; HC, healthy controls; Hcy, homocysteine; MCI, mild cognitive impairment; PD, Parkinson’s disease; SNP, single nucleotide polymorphism.

In brain cortex, subjects with dementia were also deficient in more B vitamins associated with Hcy and betaine compared to controls (**Figure 7C**): PD-D (Δ = +200%, p = 0.0008), AD-D (Δ = +131%, p = 0.030), similarly for AD-D ε4+ (Δ = +150%, p = 0.036) and ε4-(Δ = +112%, p = 0.12). PD-MCI (Δ = +58%, p = 0.26) ranked between cognitively normal and demented subjects, as if the score mirrors the cognitive status.

### Combinatorial analysis finds cognitively protective SNP combination in PD that leads to differential Hcy response to levodopa

In the last step, we combined B vitamin concentration levels with SNPs and launched a genetic programming algorithm to find combinations that could correctly assign subjects with their groups. The most successful task was aimed at distinguishing PD-D and PD-ND, resulting in identification of combined absence of rs2853533 (TYMS) and rs10380 (MTRR) as characteristic of PD-D (classification accuracy 91% for PD-D, 70% for PD-ND). 10-fold cross-validation of this approach yielded accuracies 72%, 63%, and 71% for PD-D, PD-MCI, and PD-CN, respectively. We then hypothesized that – following our results – if Hcy levels were truly implicated in dementia in PD and these SNPs were protective, we might observe their differential effects on Hcy already within PD-ND. Indeed, we found that the presence of rs2853533 or rs10380 lead to decreased Hcy levels during acute levodopa presence in PD-ND compared to subjects without these variants (**Figure 7D**). This is consistent with considering these SNPs as protective.

## Discussion

In this case-control study, we extended our previous analysis of one-carbon metabolism in AD and PD brain tissue with analysis of B vitamins and related SNPs. We found multiple associations related to cognitive impairment and dementia with multiple overlaps between PD-D and AD-D, whereas no association with PD in general was detected. Composite scores from B vitamins and SNPs associated with Hcy and betaine are consistent with their role in cognitive impairment and dementia. Importantly, we identified specific SNPs that seem to cause differential response to levodopa and would provide a mechanistic explanation for the risk of the observed levodopa-related Hcy accumulation in brain of PD-D subjects. Equally important is the finding that vitamin supplementation, as self-reported by the subjects, seem to completely ameliorate this Hcy accumulation, and could be an effective low-cost strategy to reduce the Hcy-associated cognitive decline.

We have confirmed the previous report of decreased B5 vitamin in AD^12^ and PD brain^13^ and mirrored the lower PLP in AD and PD previously reported in plasma^7,8^. In addition, we have shown that these and other B vitamin changes are related to cognitive status in PD, while non-demented groups show little differences in cortex. However, in putamen, PD-MCI subjects show similar changes as demented subjects in cortex. This might be related to the pathology progression, where substantia nigra and putamen are affected in PD first and the metabolic changes characteristic of dementia would manifest there before spreading to cortex and developing dementia.

Indeed, plasma PLP has been shown to decrease in response to chronic inflammation^33^. On the other hand, low PLP impairs Hcy clearance and there is a consensus on the causative role of Hcy in cognitive impairment^34^, accelerating the neurodegeneration. TMP is decreased similarly to PLP. Vitamins B1 and B6 metabolism is interconnected through inhibition of thiamine phosphatases by pyridoxal^35^ and regulation of pyridoxal kinase by forms of B1^36^. Increased activity of thiamine phosphatases has been previously observed in AD^37^. Therefore, it is plausible that inflammation drives lower PLP, and in turn, this causes a decrease in TMP.

The decreased pantothenic acid and biotin in demented subjects are well-correlated with PLP and support the conclusion of a more general multi-vitamin deficiency, likely resulting from a common factor. It is unclear if this could be chronic inflammation. Both pantothenic acid and biotin are transported via the sodium-dependent multivitamin transporter (SMVT) and current research suggests the opposite causal direction, where SMVT downregulation and biotin deficiency cause decreased gut permeability and inflammation^38^. Regardless of the initial trigger, pantothenic acid and biotin are crucial for energy metabolism, so their inadequate levels are detrimental and may contribute to worsening dementia.

The sudden decrease in folates in L+ PD-D coincides with Hcy accumulation and aligns with AD-D. It confirms the overall picture of high stress on the methionine cycle, where Hcy cannot be rapidly metabolized: The clearance via CBS is hampered by lower PLP, re-methylation via MTR is limited due to insufficient recovery and depletion of active folates, and re-methylation via BHMT may also be impacted: Low betaine in L-PD-D is normalized in L+ PD-D, likely reflecting the DOPA-related oxidative stress, which can affect BHMT localization and decrease its activity^39^, thereby increase betaine but decrease Hcy re-methylation.

Apart from these changes stands decreased MMA in PD-MCI putamen. While MMA is commonly regarded as an inverse indicator of vitamin B12 status, it applies specifically to adenosylcobalamin (AdoCbl), whereas Hcy re-methylation via MTR requires MeCbl. Given that we observed a trend of positive correlation between MeCbl and MMA in PD-MCI (r = 0.36, p = 0.13), the proportionality between MeCbl and AdoCbl might be inverted. Theoretically, this could happen when MTRR function is compromised and cannot re-activate oxidized cob(ii)alamin back to MeCbl, leaving cob(ii)alamin a higher chance to be converted to AdoCbl. Indeed, we detected several MTRR SNPs differentially distributed among the PD cognitive subgroups. Among them, rs1801394 shows a distinct distribution in PD-CN compared to others and is known to affect MTRR function^40^. In addition, we see that rs1801394 modulates the relationship between MeCbl and MMA in PD-MCI, leading to a trend of increased correlation with the number of mutations (0: r = −0.05, p = 0.93; Het: r = 0.37, p = 0.32; Hm: r = 0.84, p = 0.07). We thus hypothesize that the decreased MMA in PD-MCI putamen is a result of impaired MTRR function due to rs1801394 that could be of importance especially during high stress on the re-methylation pathway as with levodopa. The finding of rs1801394 Hm having increased Hcy in putamen highlights its pathological relevance.

More evidence for MTRR importance comes from rs10380 and rs162036, which are distributed differently in PD-D as if they represented a protective mutation. Unlike rs1801394, their actual effect on the protein activity has not been reported, and due to their linkage disequilibrium^10^, it is not clear if they are truly protective from dementia or if the distributions are skewed by lethality of combined mutations. Nonetheless, the opposite directions of associations with MTRR rs1801394 and rs10380/rs162036, as well as TYMS rs2853533, correspond to previous findings of associations with spina bifida^10^, where MTRR rs1801394 was protective and MTRR rs10380 and rs162036 and several TYMS SNPs were risk mutations. Spina bifida primarily stems from inadequate thymidylate levels for DNA synthesis and this part of the folate cycle can be seen as competing with the Hcy re-methylation part, as they share methylenetetrahydrofolate as an intermediate. This would explain why we saw the direction of the associations mirrored – MTRR rs1801394 as a risk mutation, while MTRR rs10380, rs162036, and TYMS rs2853533 seemed protective. TYMS rs2853533 and MTRR rs10380 were found most protective from developing dementia in PD and they led to a differential response to levodopa, which could be one of the key mechanisms of the Hcy accumulation in L+ PD-D.

The increased number of mutations in PD-MCI compared to non-demented subjects for MTR rs1805087 further implicates disturbances in Hcy re-methylation as a risk factor for cognitive impairment in PD. Another association was found for NFE2L2, which is a transcription factor that regulates the expression of various antioxidant proteins. NFE2L2 rs1806649 has been previously described as protective in PD with later age of onset^41^, although the mechanistic aspect of rs1806649 is not well-described. Our finding of higher proportion of rs1806649 mutants in PD-MCI subjects is consistent with its protective role; we saw no relation to age of onset, however, we detected a trend of increased MMSE among PD-MCI subjects (univariate linear regression β = +1.6 pt/allele, p = 0.09).

A further support for clinical implications of Hcy re-methylation disruptions comes from the associations with disease progression - BHMT rs3733890 with neurofibrillary tangles, and MTHFR rs1801131 with senile plaques. This is consistent with a previous *in vitro* study which showed that Hcy causally increases phosphorylated tau and β-amyloid peptides^42^. The opposite direction of association with MTHFR rs1801133 compared to rs1801131 and their linkage disequilibrium suggest that rs1801131 can be here more pathological than rs1801133. Cognitive impairment is known to be associated with depression, and a study in depression similarly found an association with rs1801131 and a trend (p = 0.051) of negative association with rs1801133^43^.

VNN1 rs2294757 is not well-described, with no associations reported in the literature. Our finding of link to decreased life after diagnosis, but only in demented subjects, might mean that these groups are more susceptible to disruption of pantothenic acid recycling and coenzyme A availability, which would then accelerate the disease progression. This is plausible, as experiments in a mouse model and cell cultures show that impairment of fatty acid oxidation, for which coenzyme A is required, leads to neurodegeneration with AD features^44^.

The results of associations of B vitamins with Hcy and betaine are expected, given the direct involvement of B vitamins in the one-carbon metabolism and correlations in B vitamins intake. Similarly for some associations with SNPs. However, particularly interesting is the set of associations with genes that help convert niacin to nicotinamide (BST1 rs2302465, SIRT3 rs11246020, SIRT6 rs352493). This can be explained by the subsequent methylation of nicotinamide, which generates SAH and Hcy^45^. A mutation would make the reaction less effective, generating less nicotinamide and less Hcy. This raises a question whether common vitamin supplements that contain nicotinamide might also increase Hcy in brain. In our data, the self-reported multi-vitamin intake, which might or might not contain nicotinamide, showed an overall decrease in Hcy (Welch’s t-test p = 0.028), pointing towards a full compensation by other B vitamins.

### Strengths and Limitations

We analyzed changes directly in brain tissue, which can reveal more information about neurodegeneration than surrogate measurements in biofluids. We were able to integrate measurements from one-carbon metabolism with B vitamins and genotyping data, allowing us to explore their combined effect. The samples had exceptionally low post-mortem collection intervals and were homogeneous across the groups. This is highly important for reliable comparison, as post-mortem degradation processes affect metabolism and energy balance. We were able to analyze PD subjects in relation to cognitive impairment, which revealed interesting changes. Furthermore, our analysis considered multiple confounding effects, including diseases notoriously impacting metabolism.

This study has several limitations. By design, it is an association study, which prohibits us from validating causal relationships although we discuss plausible connections with one-carbon metabolism and disease etiology. For demented subjects, we had no putamen tissue, which would help us confirm the consistency of our findings. B vitamins were analyzed without internal standards, which might lead to lower accuracy, but our QC monitoring confirmed good performance achieving low CV (mostly < 5%). Some B vitamins were not calibrated by external standards, which prevents us from calculating absolute concentrations, but this does not affect the validity of the comparisons using relative values. Despite analyzing the levodopa impact, this does not capture any potential effect of carbidopa for its substantially larger elimination halftime. Next, we did not have any information regarding diet, which would help us understand whether the changes related to B vitamins are due to inadequate intake or impaired absorption and metabolism. All subjects were non-Hispanic (where ethnicity was provided) White Americans and it is not certain whether the detected associations generalize outside this context.

### Conclusion

This study shows a widespread decrease of important B vitamins in brain tissue of demented subjects, both AD-D and PD-D. We also found genetic polymorphisms associated with cognitive impairment and dementia status in PD, while cognitively normal PD subjects often have protective genetic profiles providing them resilience against elevated Hcy and lower betaine. The groups with dementia are characterized by impairment of all three pathways for Hcy metabolism, as we previously hypothesized. These results help explain our previous finding of levodopa-associated Hcy accumulation in cortex of PD-D, and provide supportive evidence for the causative effect of levodopa-associated Hcy stress in the etiology of dementia in PD in susceptible individuals, with B vitamin supplementation as a promising therapeutic strategy.

## Acknowledgements

We are grateful to the Banner Sun Health Research Institute Brain and Body Donation Program of Sun City, Arizona, for the sale of human brain tissue. The Brain and Body Donation Program is supported by the National Institute of Neurological Disorders and Stroke (U24 NS072026 National Brain and Tissue Resource for Parkinson’s Disease and Related Disorders), the National Institute on Aging (P30 AG19610 Arizona Alzheimer’s Disease Core Center), the Arizona Department of Health Services (contract 211002, Arizona Alzheimer’s Research Center), the Arizona Biomedical Research Commission (contracts 4001, 0011, 05-901 and 1001 to the Arizona Parkinson’s Disease Consortium), and the Michael J. Fox Foundation for Parkinson’s Research.

## Funding

This study was made possible, in part, through funding received from the Aging Mind Foundation Dallas (T.B.), by the Barbara Wallace and Kelly King Charitable Foundation Trust (T.B.), and Baylor Scott & White Foundation, Dallas, Texas (T.B.).

## Author contributions

T.B. conceptualized, designed, and supervised the study. K.K. executed new metabolomic experiments. J.C.-G. and S.P. performed genotyping experiments. K.K. designed and performed data processing and statistical analysis. K.K. and T.B. contributed to the interpretation of results. K.K. wrote the initial manuscript draft. All authors (K.K., J.C.-G., S.P., T.B.) further reviewed and commented or edited the manuscript draft.

## Data availability statement

De-identified data tables with calibrated concentrations or chromatographic areas are attached as a Supplementary Data File 1.

## Disclosures and competing interests

The authors declare no competing interests.

## Notes

### Competing Interest Statement

The authors have declared no competing interest.

## References

1. Ganguly G, Chakrabarti S, Chatterjee U, Saso L. Proteinopathy, oxidative stress and mitochondrial dysfunction: cross talk in Alzheimer’s disease and Parkinson’s disease. Drug Des Devel Ther. 2017;11:797–810.

2. Boyko AA, Troyanova Nl, Kovalenko El, Sapozhnikov AM. Similarity and differences in inflammation-related characteristics of the peripheral immune system of patients with Parkinson’s and Alzheimer’s diseases. IntJ Mol Sci. 2017;18(12):2633.

3. Aarsland D, Andersen K, Larsen JP, Lolk A, Nielsen H, Kragh-Sørensen P. Risk of dementia in Parkinson’s disease: a community-based, prospective study. Neurology. 2001;56(6):730–736.

4. Kalecký K, Ashcraft P, Bottiglieri T. One-carbon metabolism in Alzheimer’s disease and Parkinson’s disease brain tissue. Nutrients. 2022;14(3):599. doi:10.3390/nu14030599

5. Seshadri S, Beiser A, Selhub J, et al. Plasma homocysteine as a risk factor for dementia and Alzheimer’s disease. N Engl J Med. 2002;346(7):476–483.

6. Folstein MF, Folstein SE, McHugh PR. “Mini-mental state”. A practical method for grading the cognitive state of patients for the clinician. J Psychiatr Res. 1975;12(3):189–198. doi:10.1016/0022-3956(75)90026-6

7. Miller JW, Green R, Mungas DM, Reed BR, Jagust WJ. Homocysteine, vitamin B6, and vascular disease in AD patients. Neurology. 2002;58(10):1471–1475.

8. Rojo-Sebastián A, González-Robles C, García de Yébenes J. Vitamin B6 deficiency in patients with Parkinson disease treated with levodopa/carbidopa. Clin Neuropharmacol. 2020;43(5):151–157.

9. Wang BJ, Liu MJ, Wang Y, et al. Association between SNPs in genes involved in folate metabolism and preterm birth risk. Genet Mol Res. 2015;14(1):850–859.

10. Shaw GM, Lu W, Zhu H, et al. 118 SNPs of folate-related genes and risks of spina bifida and conotruncal heart defects. BMC Med Genet. 2009;10:49.

11. Rajagopalan P, Jahanshad N, Stein JL, et al. Common folate gene variant, MTHFR C677T, is associated with brain structure in two independent cohorts of people with mild cognitive impairment. NeuroImage Clin. 2012;1(1):179–187.

12. Xu J, Patassini S, Begley P, et al. Cerebral deficiency of vitamin B5 (d-pantothenic acid; pantothenate) as a potentially-reversible cause of neurodegeneration and dementia in sporadic Alzheimer’s disease. Biochem Biophys Res Commun. 2020;527(3):676–681.

13. Scholefield M, Church SJ, Xu J, et al. Substantively lowered levels of pantothenic acid (vitamin B5) in several regions of the human brain in Parkinson’s disease dementia. Metabolites. 2021;11(9):569.

14. Kalecký K, Bottiglieri T. Targeted metabolomic analysis in Parkinson’s disease brain frontal cortex and putamen with relation to cognitive impairment. NPJ Parkinsons Dis. 2023;9(1):84. doi:10.1038/s41531-023-00531-y

15. Beach TG, Adler CH, Sue LI, et al. Arizona Study of Aging and Neurodegenerative Disorders and brain and Body Donation Program. Neuropathology. 2015;35(4):354–389.

16. Consensus recommendations for the postmortem diagnosis of Alzheimer’s disease. The National Institute on Aging, and Reagan Institute Working Group on Diagnostic Criteria for the Neuropathological Assessment of Alzheimer’s Disease. Neurobiol Aging. 1997;18(4 Suppl):S1-2.

17. Mirra SS, Heyman A, McKeel D, et al. The Consortium to Establish a Registry for Alzheimer’s Disease (CERAD). Part II. Standardization of the neuropathologic assessment of Alzheimer’s disease. Neurology. 1991;41(4):479–486. doi:10.1212/wnl.41.4.479

18. Movement Disorder Society Task Force on Rating Scales for Parkinson’s Disease. The Unified Parkinson’s Disease Rating Scale (UPDRS): status and recommendations. Mov Disord. 2003;18(7):738–750.

19. Adler CH, Beach TG, Zhang N, et al. Unified Staging System for Lewy Body disorders: Clinicopathologic correlations and comparison to Braak staging. J Neuropathol Exp Neurol. 2019;78(10):891–899.

20. Xu J, Clare CE, Brassington AH, Sinclair KD, Barrett DA. Comprehensive and quantitative profiling of B vitamins and related compounds in the mammalian liver. J Chromatogr B Analyt Technol Biomed Life Sci. 2020;1136(121884):121884.

21. Meisser Redeuil K, Longet K, Benét S, Munari C, Campos-Giménez E. Simultaneous quantification of 21 water soluble vitamin circulating forms in human plasma by liquid chromatography-mass spectrometry. J Chromatogr A. 2015;1422:89–98.

22. Lai SC, Nakayama Y, Sequeira JM, et al. The transcobalamin receptor knockout mouse: a model for vitamin B12 deficiency in the central nervous system. FASEBJ. 2013;27(6):2468–2475.

23. Ripley BD. The R project in statistical computing. MSOR connect 1, 23–25 (2001).

24. RStudio Team. RStudio: Integrated Development for R. (RStudio, Inc., 2019). http://www.rstudio.com

25. Fox J, Weisberg S. An R companion to applied regression. (SAGE Publications, 2018).

26. Tukey JW. Exploratory Data Analysis. (Addison-Wesley, 1977).

27. Pinheiro J, Bates DM. Mixed-Effects Models in S and S-PLUS. 1st ed. Springer; 2000.

28. Hothorn T, Hornik K, Wiel MA van de, Zeileis A. Implementing a class of permutation tests: The coin Package. J Stat Softw. 2008;28(8). doi:10.18637/jss.v028.i08

29. Warnes G, Gorjanc wcfG, Leisch F, Man M. genetics: Population Genetics [R package version 1.3.8.1.3]. CRAN (2021). https://CRAN.R-project.org/package=genetics

30. Benjamini Y, Hochberg Y. Controlling the false discovery rate: A practical and powerful approach to multiple testing. J R Stat Soc Series B Stat Methodol. 1995;57(1):289–300. doi:10.1111/j.2517-6161.1995.tb02031.x

31. Storey JD, Bass AJ, Dabney A, Robinson D. Qvalue: Q-value estimation for false discovery rate control [R package qvalue version 2.18.0]. GitHub (2019). http://github.com/idstorey/qvalue

32. Callejón G, Mayor-Olea A, Jiménez AJ, et al. Genotypes of the C677T and A1298C polymorphisms of the MTHFR gene as a cause of human spontaneous embryo loss. Hum Reprod. 2007;22(12):3249–3254.

33. Chiang EP, Smith DE, Selhub J, Da I la I G, Wang YC, Roubenoff R. Inflammation causes tissue-specific depletion of vitamin B6. Arthritis Res Ther. 2005;7(6):R1254–62.

34. Smith AD, Refsum H, Bottiglieri T, et al. Homocysteine and dementia: An international Consensus Statement. J Alzheimers Dis. 2018;62(2):561–570.

35. Lewin LM. Biosynthesis of thiamine. Inhibition of thiamine phosphatase activity by pyridoxal. Proc Soc Exp Biol Med. 1967;124(1):39–43. doi:10.3181/00379727-124-31661

36. Bunik V, Aleshin V, Nogues I, et al. Thiamine-dependent regulation of mammalian brain pyridoxal kinase in vitro and in vivo. J Neurochem. 2022;161(1):20–39. doi:10.1111/jnc.15576

37. Pan X, Sang S, Fei G, et al. Enhanced activities of blood thiamine diphosphatase and monophosphatase in Alzheimer’s disease. PLoS One. 2017;12(1):e0167273.

38. Sabui S, Bohl JA, Kapadia R, et al. Role of the sodium-dependent multivitamin transporter (SMVT) in the maintenance of intestinal mucosal integrity. Am J Physiol Gastrointest Liver Physiol. 2016;311(3):G561–70.

39. Pérez-Miguelsanz J, Vallecillo N, Garrido F, Reytor E, Pérez-Sala D, Pajares MA. Betaine homocysteine S-methyltransferase emerges as a new player of the nuclear methionine cycle. Biochim Biophys Acta. 2017;1864(7):1165–1182.

40. Olteanu H, Munson T, Banerjee R. Differences in the efficiency of reductive activation of methionine synthase and exogenous electron acceptors between the common polymorphic variants of human methionine synthase reductase. Biochemistry. 2002;41(45):13378–13385.

41. von Otter M, Bergström P, Quattrone A, et al. Genetic associations of Nrf2-encoding NFE2L2 variants with Parkinson’s disease - a multicenter study. BMC Med Genet. 2014;15(1):131.

42. Sontag E, Nunbhakdi-Craig V, Sontag JM, et al. Protein phosphatase 2A methyltransferase links homocysteine metabolism with tau and amyloid precursor protein regulation. J Neurosci. 2007;27(11):2751–2759.

43. Evinova A, Babusikova E, Straka S, Ondrejka I, Lehotsky J. Analysis of genetic polymorphisms of brain-derived neurotrophic factor and methylenetetrahydrofolate reductase in depressed patients in a Slovak (Caucasian) population. Gen Physiol Biophys. 2012;31(4):415–422.

44. Mi Y, Qi G, Vitali F, et al. Loss of fatty acid degradation by astrocytic mitochondria triggers neuroinflammation and neurodegeneration. Nat Metab. 2023;5(3):445–465. doi:10.1038/s42255-023-00756-4

45. Sun WP, Zhai MZ, Li D, et al. Comparison of the effects of nicotinic acid and nicotinamide degradation on plasma betaine and choline levels. Clin Nutr. 2017;36(4):1136–1142.

